# A Novel Syngeneic Mouse Model to Study Immune Evasion in Head and Neck Squamous Cell Carcinoma

**DOI:** 10.64898/2026.07.22.739898

**Authors:** Mari Iida, Bridget E. Crossman, Regan L. Harmon, Madeline P. Wolfe, Carlene A. Kranjac, Christine E. Glitchev, Presley N. Pergande, Rebekah M. Chacko, Sydney Van Roo, Luke W. Corday, Addison A. Sloan, Rong Hu, Peng Liu, Irene M. Ong, Haiqing Li, Prakash Kulkarni, Paul M. Harari, Justine Y. Bruce, Ravi Salgia, Deric L. Wheeler

## Abstract

Head and neck squamous cell carcinoma (HNSCC) is the sixth most common cancer worldwide, and patient outcomes have remained largely unchanged despite advances in multimodal therapy. Immune checkpoint inhibitors (ICIs), which block the PD-1/PD-L1 axis to restore T cell-mediated anti-tumor immunity, have emerged as a promising treatment strategy. However, response rates remain below 20% in HNSCC, underscoring the need to better understand mechanisms of immune evasion within the tumor microenvironment. Syngeneic mouse models are essential for studying tumor-immune interactions, yet currently available HNSCC models are limited. Here, we report the development of a novel FVB/NJ-derived syngeneic HNSCC model generated from 7,12-dimethylbenz(a)anthracene (DMBA)-induced primary on floor of mouth/buccal tumors, designated FMOC1, FMOC2, and FMOC3 (**<u>F</u>**VB/NJ **<u>M</u>**ouse **<u>O</u>**ral **<u>C</u>**ancer). *In vitro*, all FMOC cell lines exhibited robust proliferative capacity with distinct proliferation kinetics. *In vivo*, all FMOC cell lines exhibited characteristic HNSCC histopathology, including cytokeratin 5 positivity, and were tumorigenic in immunodeficient NCG mice; however, in syngeneic immunocompetent mice, only FMOC1 demonstrated sustained tumor growth at orthotopic and flank sites, whereas FMOC2 and FMOC3 tumors underwent spontaneous regression within 2 weeks, indicating differential immune-dependent tumorigenicity among the lines. Consistent with this, depletion of CD4+ and/or CD8+ T cells restored tumor growth in FMOC2 and FMOC3 models, indicating a critical role for T cell-mediated immunity in tumor suppression. Notably, FMOC1 tumors were responsive to anti-PD-L1 and anti-CTLA-4 therapy, supporting their utility for evaluating immunotherapeutic strategies. Collectively, these findings establish the FMOC model as a novel and versatile platform to study tumor-immune interactions and immune evasion mechanisms in HNSCC, with potential applications in preclinical immunotherapy development.

## Introduction

Head and neck cancer (HNC) remains a major global health challenge. HNCs arise in several anatomical sites, including the oral cavity, oropharynx, and larynx, with the majority being squamous cell carcinomas. In the United States alone, approximately 72,680 new cases of HNC are diagnosed in 2025, with over 16,680 associated deaths^1^. This data includes 59,660 cases of oral cavity and pharynx cancer and 13,020 cases of larynx cancer. Current treatment approaches for HNC include surgery, radiation, chemotherapy, and targeted therapy. Over the past decades, extensive research has identified multiple genomic alterations, including mutations in *Tp53, PIK3CA, CDKN2A* and *EGFR*^2–4^, and numerous clinical trials have evaluated targeted therapeutic strategies. Unfortunately, overall survival rates for patients with advanced disease have remained largely unchanged, with approximately 50-60% five-year survival^5^.

The introduction of immune checkpoint inhibitors (ICIs) has significantly changed the treatment landscape for recurrent or metastatic head and neck squamous cell carcinoma (HNSCC). Antibodies targeting the immune checkpoint receptor PD-1 have been approved for clinical use and have demonstrated durable responses in a subset of patients^6–10^. Nevertheless, only a minority of HNSCC patients (∼20%) benefit from immune checkpoint blockade, and both intrinsic and acquired resistance remain major clinical challenges. These observations highlight the need for a better understanding of the tumor immune microenvironment (TIME) and the mechanisms that regulate anti-tumor immune responses in HNSCC.

Many HNSCC tumors are characterized by substantial immune cell infiltration, including CD8+ cytotoxic T cells, CD4+ helper T cells, regulatory T cells (Tregs), macrophages, dendritic cells, and neutrophils^11,12^. Despite this immune presence, effective anti-tumor immunity is often suppressed within the TIME. One major mechanism involves inhibitory immune checkpoint signaling through pathways such as PD-1/PD-L1, which dampens cytotoxic T cell activity. In addition, immunosuppressive cell populations are frequently enriched in HNSCC tumors. Tregs inhibit cytotoxic T cell activation and produce anti-inflammatory cytokines, while tumor-associated macrophages (TAMs) often adopt an M2-like phenotype that promotes tumor growth, angiogenesis, and immune suppression. The accumulation of myeloid-derived suppressor cells (MDSCs) further suppresses T cell activation and contributes to immune tolerance. Tumor-derived cytokines such as IL-10, TGF-b and VEGF also play critical roles in shaping this immunosuppressive TIME.

Preclinical syngeneic mouse models are essential tools for investigating tumor-immune interactions and evaluating immunotherapeutic strategies. Several models have been developed for HNSCC research. The Mouse Oral Cancer (MOC) series (MOC1, MOC2, and MOC22) was generated from 7,12-Dimethylbenz[a]anthracene (DMBA)-induced oral tumors in C57BL/6 mice^13–16^. Previous studies, including work from our laboratory, have demonstrated that MOC1 tumors are moderately immunogenic and exhibit higher CD8+ T cell infiltration, whereas MOC2 tumors are poorly immunogenic and resistant to immune checkpoint blockades^16–18^. Another model system, the 4MOSC series (4MOSC1 and 4MOSC2), was derived from 4-nitroquinoline-1-oxide (4NQO) induction in C57BL/6 mice^19^. In this system, 4MOSC1 tumors display moderate immunogenicity and partial sensitivity to anti-PD-1 therapy, while 4MOSC2 tumors are highly aggressive and resistant to checkpoint inhibition. In addition, the SCC7 cell line, derived from spontaneous squamous cell carcinoma in C3H mice, has been widely used in studies examining radiation-immunology combinations^20,21^. Despite these advances, the number of available syngeneic mouse models for studying tumor-immune interactions in HNSCC remains limited compared with the large number of established human HNSCC cell lines.

To address this limitation, we developed a novel syngeneic HNSCC model in FVB/NJ mice derived from primary on the floor of mouth/buccal tumors induced by DMBA. After 26 weeks of carcinogen treatment, we established three FVB/NJ mouse oral cancer cell lines, designated FMOC1, FMOC2, and FMOC3. FMOC1 tumors grew rapidly and exhibited metastatic potential, whereas FMOC2 and FMOC3 tumors failed to grow or metastasize under normal conditions. Notably, T-cell depletion restored tumor growth in FMOC2 and FMOC3 models, highlighting a critical role for adaptive immunity in controlling these tumors. Furthermore, FMOC1 tumors were sensitive to treatment with anti-PD-L1 or anti-CTLA-4 therapeutic antibodies. These newly established syngeneic models in the FVB/NJ mouse background provide additional tools for investigating tumor-immune interactions and may contribute to the development of improved immunotherapeutic strategies for HNSCC.

## Materials and Methods

### Establishment of cell lines and tissue culture

Animal procedures and maintenance were conducted in accordance with the University of Wisconsin — Madison School of Medicine and Public Health IACUC guidelines. Six male and eight female FVB/NJ mice (5–6 weeks of age) were purchased from UW Biomedical Research Model Services (Madison, WI, USA). Mice were treated with 25 μg of 7,12-dimethylbenz(a)anthracene (DMBA, Sigma) dissolved in 20 μL of ethanol, administered by oral gavage twice weekly. Mice were euthanized when institutional criteria for experimental neoplasia were met. Following euthanasia, mice were photographed and complete necropsies were performed. Tumor specimens were collected, and portions were either flash-frozen in liquid nitrogen or fixed in 10% neutral-buffered formalin for 24 hours at 4C. Individual lesions were dissected, and tumor cells were isolated to establish FMOC cell lines.

### Immunoblot analysis

Whole-cell protein lysates were prepared using RIPA buffer (50 mM HEPES, pH 7.4; 150 mM NaCl; 0.1% Tween-20; 10% glycerol; 2.5 mM EGTA; 1 mM EDTA; 1 mM DTT; 1 mM Na_3_VO_4_; 1 mM PMSF; 1 mM b-glycerophosphate; and 10 μg/ml leupeptin and aprotinin). Immunoblotting was performed as previously described ^22–26^. Antibodies were used according to the manufacturer’s instructions: Cyclin D1 (Cell Signaling Technologies, MA, USA, #55506), GM-CSF (Proteintech, Inc., IL, USA #17762-1-AP), GAPDH (CST #2118), α-tubulin (Proteintech, Inc #66031-1-Ig).

### Cell proliferation assay and BrdU flow cytometry assay

Cell proliferation assays were performed using FMOC1, FMOC2, and FMOC3 cells. Cells were seeded on a 96-well plate and stained with Incucyte Nuclight Rapid Red Dye (Sartorius 4717, Göttingen, Germany). To monitor cell proliferation over time, the plate was placed in the Incucyte Live Cell Imaging System (Sartorius) and images were acquired every 4 hours. Data analysis was performed using a red fluorescence mask to accurately count each cell nucleus over time in the Incucyte software.

For BrdU analysis, cells were plated at 300,000 cells per 100-mm dish and allowed to adhere overnight. Cells were pulsed with 10 μM BrdU for 1 h, harvested using Accutase (Invitrogen #004555-56), and stained with an APC-conjugated anti-BrdU antibody according to the manufacturer’s instructions (BD Pharmingen, Franklin Lakes, NJ, #552598). Samples were analyzed using an Attune Nxt flow cytometer (Thermo Fisher Scientific, Waltham, MA, USA), and data were processed using FlowJo Software (BD, RRID: SCR_008520).

### Wound healing assay (Gap Closure assay)

FMOC cells (5 × 10⁴ cells/70 μL per well) were seeded into Culture-Insert 2 well chambers (Ibidi, Munich, Germany) and incubated overnight. Inserts were removed to create a uniform gap. Migration was measured 0, 2, 5, 7, 9, 12, and 24 hours using an inverted microscope. Migration was quantified by measuring gap width at three positions, and migration rate was calculated as: [(initial gap width - final gap width) / initial gap width] × 100.

### DNA Isolation, Whole-Exome sequence analysis

Genomic DNA was extracted using the DNeasy Blood & Tissue Kit (Qiagen). Whole-exome libraries were prepared using the Illumina Genomic DNA library prep kit and enriched using the Twist Mouse Exome kit performed by UW Biotechnology core. Captured libraries were sequenced on an Illumina NovaSeq 6000. Illumina’s DRAGEN software (version 3.10) was used to align reads to mouse genome (mm10) and call variants. Functional impact of variants was predicted by Ensembl’s Variant Effect Predictor (cache version 102).

### RNA Isolation, cDNA Synthesis, and qPCR

RNA isolation, cDNA synthesis, and qPCR were performed as previously described ^23–25^. Gene expression was measured using TaqMan Fast Advanced Master Mix and Probes (Thermo Fisher Scientific). Relative expression was calculated using the ΔΔCT method, with GAPDH and ActinB as internal controls.

### In Vivo Mouse Experiments

All procedures were approved by the University of Wisconsin — Madison IACUC guidelines. Cells were injected subcutaneously into the dorsal flank of 4-6-week-old female FVB/NJ mice (Jackson Laboratory, Bar Harbor, ME, USA), athymic nude (Envigo, Indianapolis, IN, USA), or NCG (Charles River Laboratories, Wilmington, MA, USA) mice. Tumors were measured two times per week using a digital caliper. For orthotopic experiments, the cells were injected to the right buccal mucosa area via the intraoral route at a final concentration of 1 x10^6^/0.1 ml per animal. Cells were resuspended in PBS containing 50% Cultrex RGF BME Matrigel (Corning Inc., Corning, NY, USA, #3433-005-01) prior to inoculation. For metastasis potential experiments, single-cell suspensions of tumor cells were prepared prior to tail vein injection. Mice were intravenously (IV) injected with 1 × 10^6^ cancer cells in 100 μL PBS in the tail vein using insulin syringes.

### Tumor Dissociation and Flow Cytometry

Tumors were harvested and dissociated as previously described ^18,27^. Cells were stained with viability dye, blocked with Fc receptor blocker (Biolegend, SanDiego, CA, USA), and stained with surface antibodies. Intracellular staining was performed using FoxP3/Transcription Factor Staining Buffer Kit (Tonbo Biosciences, San Diego, CA, USA). Samples were analyzed using an Attune NxT flow cytometer and FlowJo software (BD, RRID: SCR_008520). Antibodies and gating strategies are listed in Supplemental Table S2 and S3.

### T cell-killing Assay and Annexin V Analysis

Splenocytes were isolated from tumor-bearing FVB/NJ mice and CD8+ T cells were purified using the MojoSort Mouse CD8^+^ T Cell Isolation kit (Biolegend). For cytotoxicity assays, 5 × 10^3^ FMOC cells were co-cultured with 5 × 10^4^ CD8^+^ T cells in 96-well plate. After 4 h at 37^◦^C, Annexin V-FITC was added, and fluorescence was measured using a plate reader.

### T-cell Depletion

CD4^+^ and/or CD8^+^ T cells were depleted using anti-CD4 (clone GK1.5) and anti-CD8α (clone 2.43) antibodies (BioXcell, Lebanon, NH). Mice received 300 µg intraperitoneally on days −5 and 0, followed by maintenance dosing every 5 days. Depletion efficiency (>90%) was confirmed by flow cytometry.

### Immunohistochemistry (IHC)

Tumors were fixed in 10% neutral buffered formalin, paraffin-embedded, and sectioned. Antigen retrieval was performed using 10 mM citrate (pH 6.0) or 10 mM Tris-1 mM EDTA (pH 9.0) buffer. Sections were stained using antibodies (Supplemental Table S1) and detected using either the ImmPRESS HRP Goat Anti-Rat or Goat Anti-Rabbit IgG Polymer Detection Kit (Vector Laboratories, Burlingame, CA, USA) in conjunction with 3,3’-diaminobenzidine substrates (Vector Laboratories). Slides were counterstained with hematoxylin (Thermo Fisher Scientific) and imaged using an Olympus BX51 microscope. Quantitation of staining intensity was performed using FIJI version 1.54 ^17,18,27^.

### Immunofluorescence

Cells were plated in 4-well Millicell EZ chamber slides (Millipore Sigma, Burlington, MA, USA) for 24 hours and fixed in 3% formaldehyde, permeabilized in 0.3% Triton X-100, and blocked in normal goat serum. Cells were stained with cytokeratin 5 primary antibody (Proteintech, 1:100, 28506-1-AP) overnight at 4^◦^C, followed by goat anti-rabbit Alexa Fluor 546 secondary antibody (Invitrogen). Actin was stained using the ActinGreen 488 ReadyProbes Reagent (Invitrogen), and nuclei were stained with DAPI. Imaging was performed using a Nikon AXR Confocal Microscope.

### Cytokine Array

Cells were plated (50,000 cells/mL; 1mL) in serum-free media and incubated for 48hr. Supernatant was collected and cytokine and chemokine levels from cell culture supernatant were measured using the Mouse Cytokine Array 3 (RayBiotech, Peachtree Corners, Georgia, USA, AAM-CYT-3-2) according to manufacturer’s instructions. Membranes were imaged on Azure 600 imager (Azure Biosystems) and analyzed using FIJI version 1.54.

### Luminex Assay

Cells were plated (250,000 cells/mL; 2mL) in serum-free media and incubated for 24 hours. Supernatants were collected and cytokine and chemokine levels from cell culture supernatant were measured using the ProcartaPlex Mouse Cytokine and Chemokine Panel 1, 26 plex (Thermo Fisher Scientific, EPX260-26088-901) according to manufacturer’s instructions. Data were acquired using a Luminex xMAP INTELLIFLEX System (Thermo Fisher Scientific) and analyzed with ProcartaPlex software (Thermo Fisher Scientific).

### Bulk RNA Sequencing

RNA was extracted from FMOC1, FMOC2, and FMOC3 cells using the RNeasy Plus Mini kit (Qiagen, Hilden, Germany). The sample libraries were sequenced on an Illumina HiSeq4000 (Illumina, San Diego, CA) to generate 101 bp paired-end reads. Raw RNA-seq reads were processed by trimming adaptor and poly(A) sequences using fastp (v0.23.2)^28^. The cleaned reads were aligned to the mouse reference genome (mm10) using STAR (v2.7.9) ^29^ with gene annotations from GENCODE v38^30^. Gene-level expression counts were quantified using the featureCounts function from the Subread package (v2.0.3)^31^. Genes with very low expression (maximum expression <0.1 RPKM across all samples) were excluded from downstream analysis. Differential expression analysis was performed on raw count data using DESeq2 (v1.38.2)^32^, and differentially expressed genes (DEGs) were identified using a false discovery rate (FDR) threshold of <0.05. Heatmaps of DEGs were generated using the R pheatmap package (v1.0.12)^33^, while volcano plots and dot plots were created using ggplot2 (v3.4)^34^. Overlapping DEGs were visualized using Venn diagrams generated in R ggvenn (v0.1.19)^35^. The circular heatmap was generated using the circlize (v0.4.17)^36^ and ComplexHeatmap (v2.25.2)^37^ packages in R. Pathway-level expression values across the three FMOC cell lines (FMOC1, FMOC2, and FMOC3) were row-scaled (z-scored) to enable comparison of relative expression patterns.

### Statistical analysis

Statistical significance was defined as P<0.05. Analysis of flow cytometry, immunohistochemistry, and Luminex was performed using GraphPad Prism (GraphPad Software, Inc; RRID: SCR_002798). Two-group comparisons used two-sided t-tests, while multiple groups were analyzed using one-way ANOVA with Dunnett’s post hoc tests.

### Data availability

The whole exome sequencing data of murine DMBA-induced syngeneic cell lines have been deposited in the NCBI GEO accession (GSE332802, reviewer token ‘qbuveukinbmvjmb’). The source data underlying Fig. 1F are provided as Supplementary Data in Microsoft excel format. All the other data supporting the findings of this study are available within the article and its Supplementary Information files and from the corresponding author upon reasonable request.

**Figure 1.**
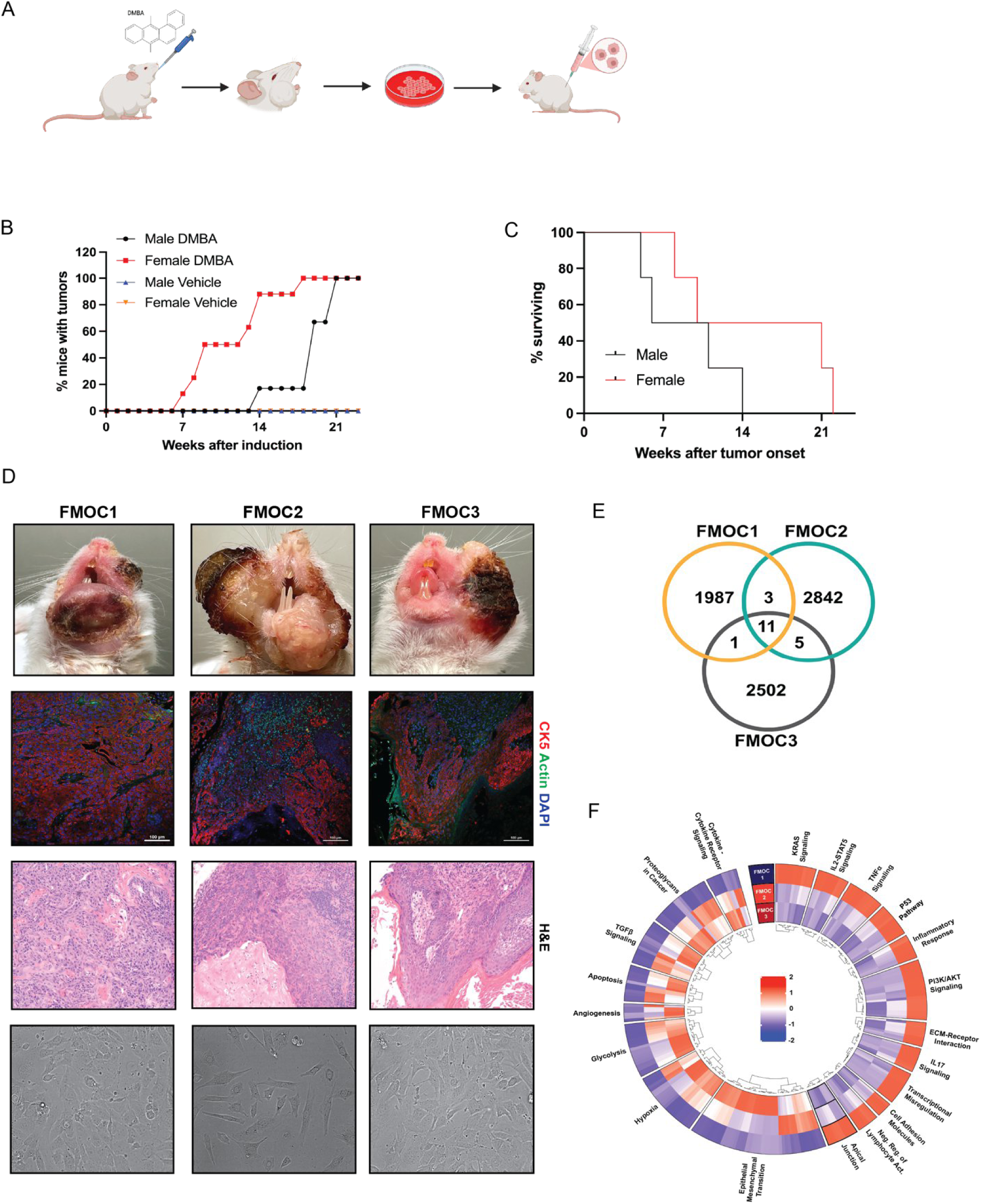
Development of a novel FVB/NJ-derived syngeneic HNSCC mouse model. (A) Experimental scheme for developing the FMOC syngeneic model. (B) The percentage of FVB/NJ mice with tumors after DMBA or vehicle treatment. (C) The percentage of surviving FVB/NJ mice after tumors developed. (D) Top panel; Generated 3 FMOC cell lines from independent DMBA-induced tumors. Upper-middle panel; Immunofluorescent staining of cytokeratin 5 (CK5, red), Actin (green) and DAPI (blue) to show squamous cell characterization of the lesion in each mouse with FMOC tumors (bar 100um). Lower-middle panel, representative H&E-stained section of primary tissue from each FMOC tumor (200X magnification). Bottom panel, representative 4x microscopic image of FMOC cell lines by Evos XL Core. (E) Overlap of high or moderate variants from whole exome sequences by Venn diagram. (F) General heatmap for pathways with significant differential gene expression among FMOC cell lines by bulk-RNA sequencing analysis.

## Results

### A novel DMBA-induced oral squamous cell carcinoma model in FVB/NJ mice

To establish a murine model for studying immune evasion and suppression, male (n=6) and female (n=8) FVB/NJ mice were treated with 7,12-dimethylbenz(a)anthracene (DMBA) twice weekly (**Figure 1A**) for up to 26 weeks. DMBA is a potent polycyclic aromatic hydrocarbon (PAH) and a well-established chemical carcinogen widely used in experimental cancer models including oral cavity squamous cell carcinoma^14^. Both male and female mice developed visible nodules on the floor of mouth or buccal mucosa between 6 and 13 weeks after treatment initiation (**Figure 1B**), and 100% of mice progressed to tumor formation. In contrast, no tumors were observed in vehicle-treated controls (n = 10) over the 26-week period. Male mice reached euthanasia criteria more rapidly than females (**Figure 1C**). All tumor-bearing males were euthanized within 14 weeks of tumor onset, whereas all female mice reached endpoint by 22 weeks. From 14 independent DMBA-induced tumors, we established three transplantable mouse oral cancer cell lines, designated FVB/NJ mouse oral cancer 1, 2, or 3 (FMOC1, FMOC2, and FMOC3; **Figure 1D**). Histological analysis confirmed features consistent with HNSCC histology, as demonstrated by cytokeratin 5 immunofluorescence, hematoxylin and eosin (H&E) staining and microscopic image of FMOC cell lines. To characterize the mutational landscape of these cell lines, we performed whole-exome sequencing. Approximately 2,000-3,000 variants were identified per sample, with minimal overlap among the three lines, indicating distinct mutational profiles (**Figure 1E**, NCBI GEO accession: GSE332802). The predominant base substitution types were T>A and C>A transversions (**Supplemental Table S4**), consistent with DMBA-induced mutagenesis and carcinogen-associated mutational patterns reported in subsets of HNSCC ^3^. Each FMOC line harbored mutations in key oncogenes and tumor suppressor genes. FMOC1 exhibited alterations in *KRAS, MYCN, BCL2, FGFR2, MET, RET, ROS1, ABL1, TNK2, TP53,* and *DLC1*. FMOC2 contained mutations in *FGFR1, HRAS, NOTCH1, TP53, APC,* and *NF1*. FMOC3 harbored mutations in *EGFR, HER3, MET, PIK3CA,* and *TP53*. Furthermore, bulk-RNA sequencing analysis confirmed exome analysis: Kras, IL-2/STAT5, TNFa, and PI3K/AKT gene sets were upregulated in FMOC1 cells, whereas TGFβ signaling or apoptosis pathway genes were upregulated in FMOC2 and FMOC3 cells (**Figure 1F and Supplemental Table S5**). Collectively, these data demonstrated that FMOC tumors exhibit alterations in receptor tyrosine kinase (RTK), PI3K, and TP53 signaling pathways, consistent with previously reported DMBA-induced oral cancer models such as the MOC series ^15,16^. These findings establish FMOC1, FMOC2, and FMOC3 as novel syngeneic mouse models of oral squamous cell carcinoma.

### Cell proliferation potential in FMOC oral squamous cell model

During in vitro cell culture, we observed differences in the growth rates of FMOC1, FMOC2 and FMOC3 cell lines. To assess proliferative capacity, we performed real-time quantification of cell proliferation using the Incucyte Live-Cell Analysis System (**Figure 2A**). The proliferation curve showed that FMOC1 cells grow rapidly after 48 hours whereas FMOC2 and FMOC3 cells grow moderately. These results indicate that all FMOC cell lines are capable of in vitro growth, with distinct proliferation kinetics. Next, we evaluated cell motility using a wound healing assay. After 12h following removal of the insert, FMOC1 demonstrated rapid gap closure (∼80% closure), whereas FMOC2 and FMOC3 exhibited slower migration (∼30-40% closure) (**Figure 2B and Supplemental Figure 1**). To further characterize proliferative activity, we performed a BrdU incorporation assay analyzed by flow cytometry. The distribution of cells in S-phase and the extent of BrdU incorporation differed among the FMOC lines (**Figure 2C**). Notably, FMOC1 displayed increased early S-phase accumulation compared to FMOC2 and FMOC3. Given these differences, we examined expression of one of the cell cycle regulators, cyclin D1 by immunohistochemistry (**Figure 2D**) and immunoblot analysis (**Figure 2E**). Cyclin D1 expression was markedly higher in FMOC1 compared to FMOC2 and FMOC3. Collectively, these data demonstrated that while all FMOC cell lines proliferate *in vitro*, FMOC1 exhibits a more aggressive phenotype characterized by faster proliferation, enhanced migratory capacity, and increased cell cycle activity relative to FMOC2 and FMOC3.

**Figure 2.**
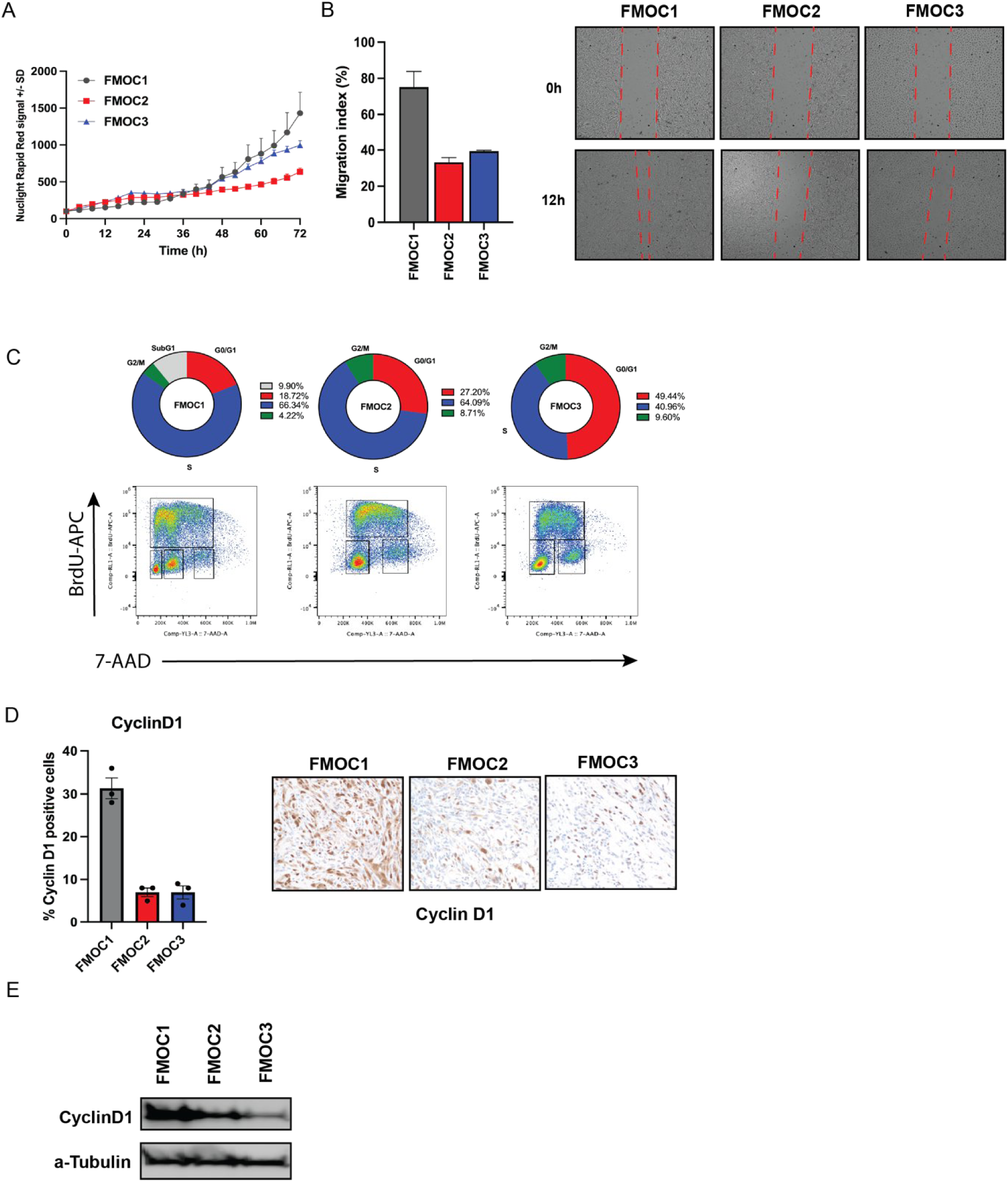
Differential proliferation, migration, and cell cycle dynamics among FMOC cell lines. (A) FMOC cell proliferation plot by Incucyte (n=6). (B) Wound healing assay of FMOC cell lines after 12h incubation. Mean values and SEMs are shown. (n = 3 per photo). (C) BrdU incorporation assay. (n=3 per cell line) (D) Cyclin D1 expression levels in FMOC tumors by IHC (n = 3 tumors per group). Representative images of Cyclin D1 staining in FMOC tumors are shown (400X magnification). (E) Whole cell lysate was collected from FMOC cell lines and immunoblotted for Cyclin D1. α-tubulin was used as a loading control.

### Immunosuppressive properties of the tumor microenvironment vary among FMOC cell lines

To evaluate *in vivo* tumor growth, FMOC cell lines were inoculated bilaterally into the flanks of NCG, athymic nude or syngeneic FVB/NJ mice via subcutaneous injection, and tumor growth was monitored for up to 70 days. All three FMOC cell lines successfully formed tumors in highly immunodeficient NCG mice, which lack T cell, B cell and natural killer (NK) cells (**Figure 3A**), as well as in athymic nude mice, which lack T cells (**Figure 3B**). In both models, FMOC1 tumors exhibited the most rapid growth, followed by FMOC2 and FMOC3. Strikingly, in syngeneic FVB/NJ mice, FMOC2 and FMOC3 initially formed tumors that spontaneously regressed after approximately 2 weeks, whereas FMOC1 tumors continued to grow progressively (**Figure 3C**). To assess tumor growth at the primary anatomical site, FMOC cell lines were orthotopically implanted into the floor of the buccal region (n=8 per group) in athymic nude (**Figure 3D**) or syngeneic FVB/NJ (**Figure 3E**) mice. In athymic nude mice, all FMOC1 (8/8), FMOC2 (8/8) and FMOC3 (8/8) tumors successfully established growth (**Figure 3D**), whereas all FMOC1 tumors (8/8) and only one FMOC2 tumor (1/8) and none of the FMOC3 tumors (0/8) formed tumors in FVB/NJ mice (**Figure 3E**). Given the high propensity of HNSCC for locoregional lymph node metastasis and its association with poor prognosis, we next evaluated metastatic potential using a tail vein injection model in both athymic nude and syngeneic FVB/NJ mice. In athymic nude mice, all FMOC1 (9/9), and FMOC3 (8/8) tumors metastasized to the lung, pleura or heart (**Figure 3F**). One FMOC3 mouse developed a large left submandibular mass and enlarged lymph nodes (data not shown). Interestingly, none of the FMOC2 tumors (0/8) metastasized to the lung (**Figure 3F**). FVB/NJ mice injected with FMOC1 cells developed metastatic lung colonization, with 73% (11/15) exhibiting lung nodules. In contrast, no lung metastases were observed in FVB/NJ mice injected with FMOC2 or FMOC3 cells (0/15 for both) (**Figure 3G**). Collectively, these results suggest that differences in tumor growth among FMOC cell lines are influenced by immune-dependent mechanisms, with FMOC1 potentially exhibiting a more immunosuppressive TIME that supports tumor progression and metastasis.

**Figure 3.**
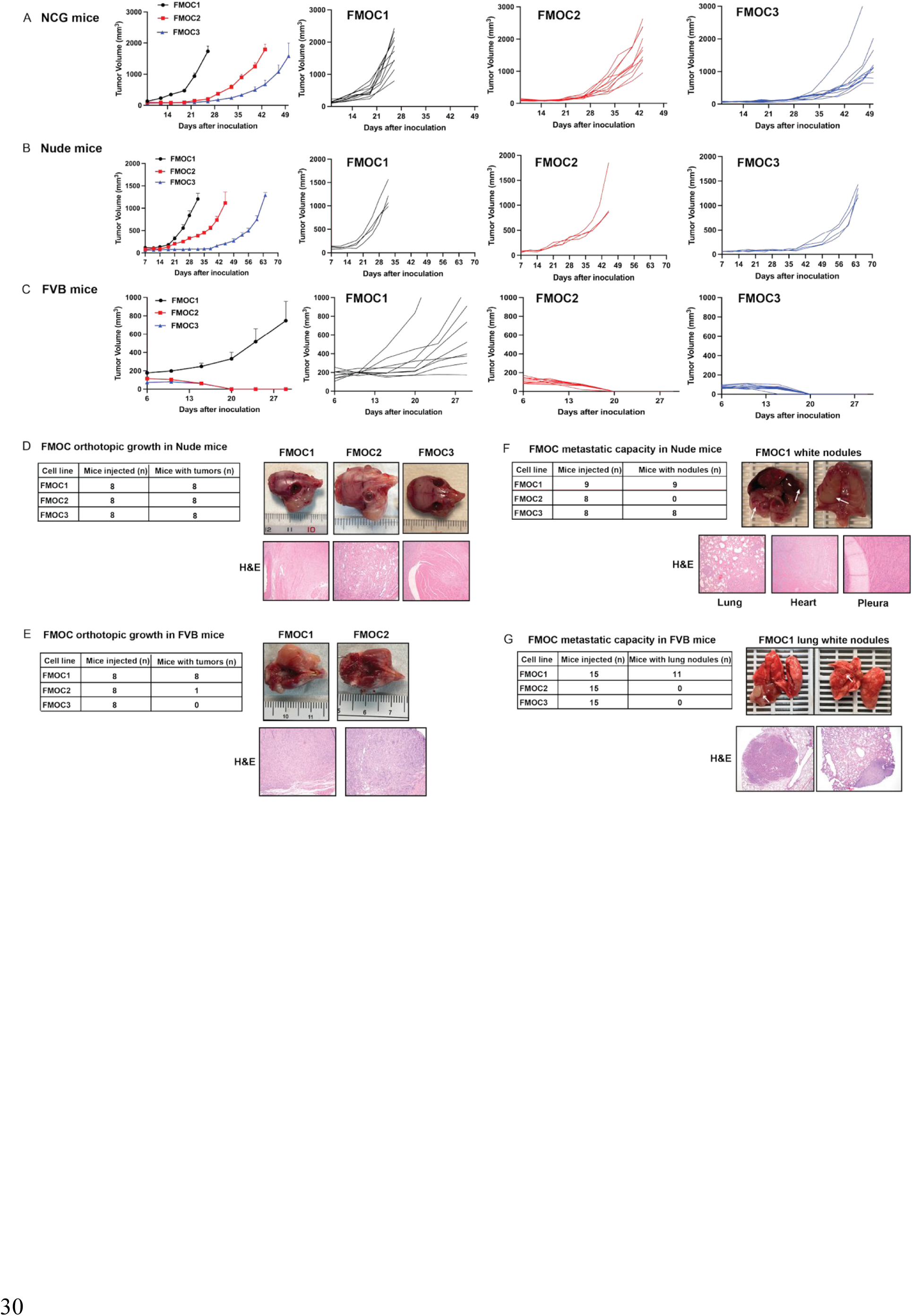
Immunosuppressive properties of the tumor microenvironment vary among FMOC cell lines. (A-C) 1× 10^6^ cells/tumor were subcutaneously injected in NCG (A), athymic nude (B) or syngeneic FVB/NJ (C) mice. Tumor volume was measured twice per week. Mean values and SEMs are shown (n = 4-10 tumors per group). Individual tumor growth rates are shown. (D, E) Summary of orthotopic tumor growth of each cell line in athymic nude (D) and FVB/NJ (E) mice. Representative H&E-stained section of tumor showing moderately differentiated squamous cell carcinoma (100X, 200X magnification). (F, G) summary of metastatic capacity in lung, heart or pleura of each cell line in athymic nude (F) and FVB/NJ (G) mice. White arrows indicated nodules in lung, heart or pleura. H&E-stained metastatic nodule shows effacement of normal architecture by squamous cell carcinoma (40X, 100X or 200X magnification). H&E; hematoxylin and eosin stain.

### Tumor growth in the FMOC series is associated with distinct immune cell populations

Given the observed differences in FMOC1, FMOC2, and FMOC3 tumor growth across mouse models with variable immune constitutions, we next characterized the immune cell composition within the TIME. We performed flow cytometry to assess levels of tumor-infiltrating leukocytes (TILs) in FMOC tumors at 10 days post-inoculation onto the flanks of syngeneic FVB/NJ mice. Overall, FMOC2 and FMOC3 tumors exhibited high levels of anti-tumor immune infiltration, including increased frequencies of CD8^+^ T cells, cytotoxic CD8^+^ T cells, and CD4^+^ T cells, natural killer T (NKT) cells and Basophils, as well as enhanced activation and cytotoxicity of natural killer (NK) cells. Further, we observed enrichment of M1 macrophages and classical dendritic cells (DCs) in FMOC2 and FMOC3 tumors (**Figure 4A**). In contrast, FMOC1 tumors showed enrichment of immunosuppressive cell populations, including regulatory T cells (Tregs) and M2 macrophages (**Figure 4B**). FMOC1 tumors were also enriched for cell populations that can have both anti-and pro-inflammatory effects, including neutrophils and eosinophils (**Figure 4B**). Immunohistochemical analysis further supported these findings, demonstrating higher levels of CD8, CD4, F4/80 and NK1.1 staining in FMOC2 and FMOC3 tumor compared to FMOC1 (**Figure 4C**). While FoxP3 staining was similar across the three models, the CD8/FoxP3 ratio was highest in FMOC3 tumors. Collectively, these results indicated that tumor growth differences among FMOC cell lines are associated with distinct immune microenvironments, with FMOC2 and FMOC3 characterized by enhanced anti-tumor immunity, whereas FMOC1 exhibits a more immunosuppressive TIME.

**Figure 4.**
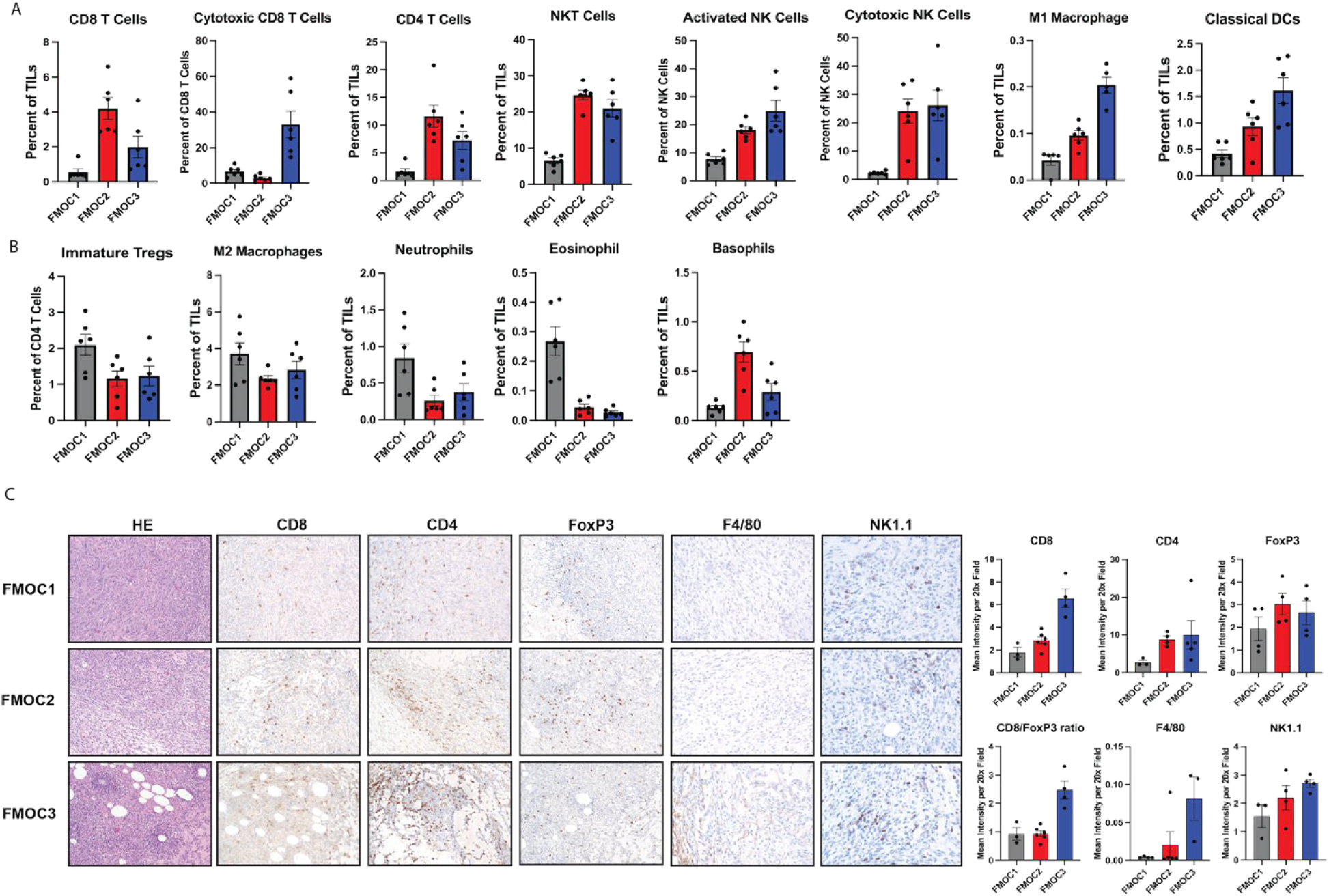
Tumor growth in the FMOC series is associated with distinct immune cell populations. (A, B) Subcutaneous tumors were generated in FVB/NJ mice by inoculating 1 × 10^6^ of each FMOC cells per tumor. Tumors were collected 10 days after inoculation, and tumor-infiltrating lymphocytes (TILs) were analyzed via flow cytometry. Mean values and SEMs are shown (n = 5-6 tumors per group). (C) Representative images of H&E and Tumor immune infiltrate (CD8, CD4, FoP3, F4/80, and NK1.1) analyzed by IHC for each cell line are shown (200X magnification). Image quantification was performed, and the CD8α/FoxP3 ratio was calculated. Mean values and SEMs are shown (n = 3-6 images per group).

### T cells contribute to tumor control in FMOC2 and FMOC3 models

Because FMOC2 and FMOC3 tumors failed to grow in syngeneic FVB/NJ mice and exhibited high infiltration of CD8^+^ and CD4^+^ T cells compared to FMOC1, we hypothesized that T cell-mediated immunity restricts tumor growth in these models. To test this, FVB/NJ mice were treated with CD4^+^ and CD8^+^ T-cell-depleting antibodies prior to, and throughout, subcutaneous inoculation with FMOC2 or FMOC3 cells. Efficient depletion was confirmed by flow cytometry following intraperitoneal administration of anti-CD8 and anti-CD4 antibodies at day 43 (**Figure 5A and 5B**). Notably, combined depletion of both CD8^+^ and CD4^+^ T cells restored tumor growth in both FMOC2 and FMOC3 models compared to the IgG-treated control group (**Figure 5C and 5D**). To further dissect the contribution of individual T cell subsets, we depleted CD8^+^ or CD4^+^ T cells separately. Depletion efficiency was confirmed at day 41 (**Figure 5E and 5F**). In FMOC2 tumors, depletion of either CD8^+^ or CD4^+^ cells alone was sufficient to permit tumor growth in FVB/NJ mice (**Figure 5G**). In contrast, FMOC3 tumors required depletion of both CD8^+^ and CD4^+^ T cells to achieve robust growth, suggesting a cooperative role for these subsets in tumor control (**Figure 5H**).

**Figure 5.**
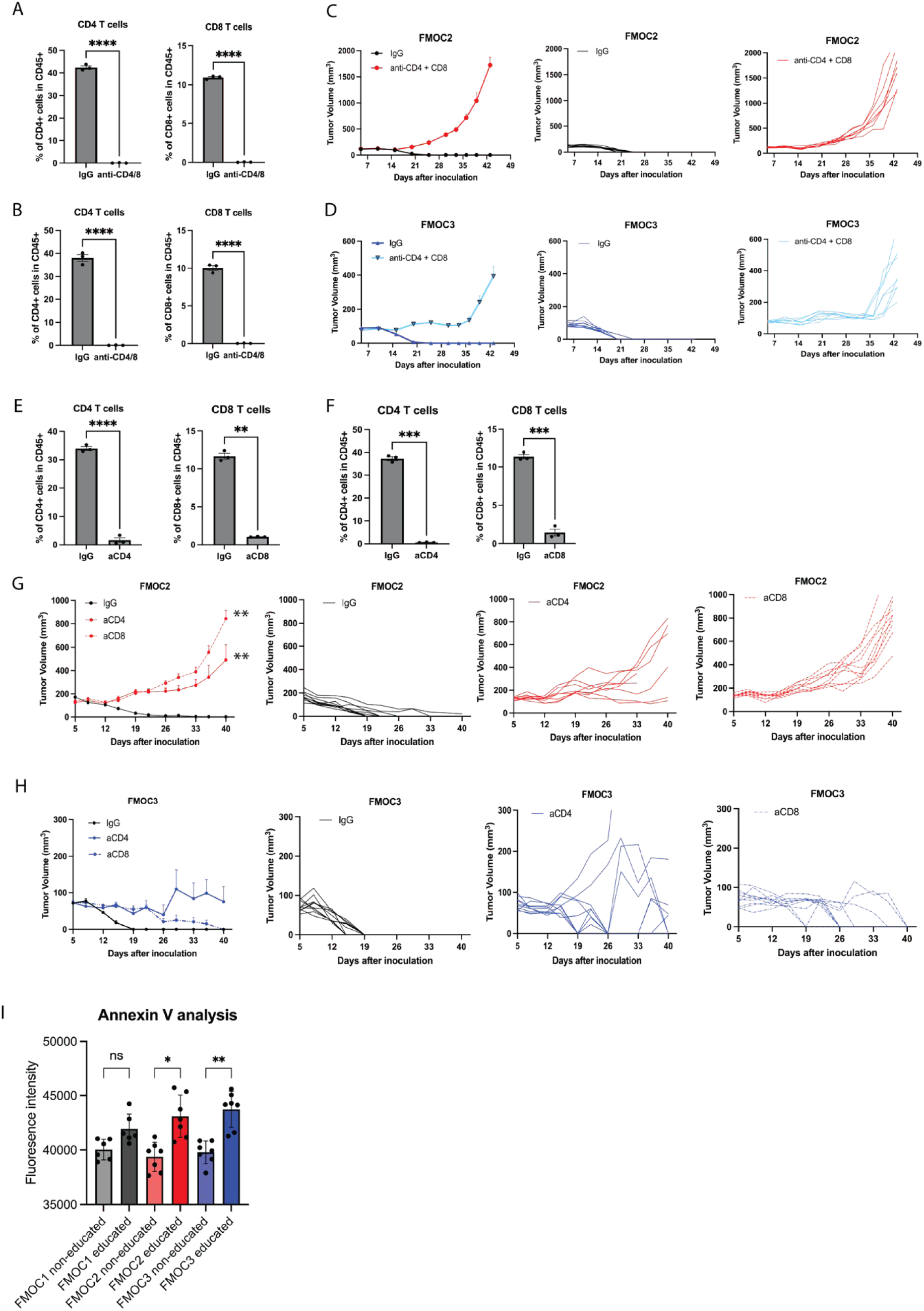
T cells contribute to tumor control in FMOC2 and FMOC3 models. (A, B) Spleens were collected to confirm the simultaneous depletion of CD4+ and CD8+ cells in FMOC2 (A) or FMOC3 (B) bearing mice at day 43 (n = 3 spleens per group). (C, D) The combination of anti-CD4 and anti-CD8 treatment restore tumor growth in FMOC2 (C) and FMOC3 (D) mice. Individual tumor growth rates are shown. (n = 7-10/group). (E, F) Spleens were collected to confirm the selective depletion of either CD4+ or CD8+ cells in FMOC2 (E) or FMOC3 (F) bearing mice at day 41 (n = 3 spleens per group). (G) FMOC2 tumor growth was rescued by CD4 or CD8 simultaneous depletion. Individual tumor growth rates are shown. (n = 8-10/group) (H) FMOC3 tumor growth was not restored in FVB/NJ mice depleted of CD4 or CD8 T cells. Individual tumor growth rates are shown. (n = 8-10/group) (I) Education toward FMOC2 or FMOC3 enhances T cell cytotoxicity. Primary CD8+ cells (5 × 10^4^ per well) were cultured with each FMOC cells (5 × 10^3^ per well) for 4 h. Annexin V was measured, and fluorescent intensity was calculated. ns, not significant; * p < 0.05; ** p < 0.01.

To evaluate the cytotoxic potential of T cells, we performed *in vitro* co-culture assays using primary CD8^+^ T cells isolated from tumor-bearing (“educated”) or naïve (“non-educated”) mice. Annexin V analysis demonstrated that educated CD8^+^ T cells induced significantly higher levels of apoptosis in FMOC2 and FMOC3 cells compared to non-educated controls (**Figure 5I**). In contrast, no significant difference in apoptosis was observed in FMOC1 cells. These results provide functional evidence that immune-mediated tumor rejection in FMOC2 and FMOC3 is primarily driven by T cell activity. Collectively, these findings demonstrate that T cell-mediated immunity plays a critical role in controlling FMOC2 and FMOC3 tumor growth, whereas FMOC1 appears resistant to T cell-mediated cytotoxicity, consistent with its immunosuppressive TIME.

### Response of FMOC1 tumors to immune checkpoint inhibitors

Given the bulk RNA-seq data that indicated modulation of cytokine and cytokine receptor signaling pathways, we next explored cytokine and chemokine expression profiles of FMOC cell lines. To accomplish this, we utilized a Mouse Cytokine Array (RayBiotech, Peachtree Corners, GA). This cytokine array identified several key immunomodulatory factors in FMOC cell lines, including granulocyte-macrophage colony-stimulating factor (GM-CSF), IFN-g, IL-10, sTNFR1, IL-17 and CCL5 (**Figure 6A**). High GM-CSF levels were confirmed in FMOC1 cells by qPCR (**Figure 6B**), Luminex ProcartaPlex Mouse Cytokine and Chemokine Panel 1 (Invitrogen) analysis (**Figure 6C**) and immunoblotting (**Figure 6D**). Given the role of GM-CSF in modulating myeloid cell activity and antigen presentation, which can influence responses to PD-1/PD-L1 and CTLA-4 checkpoint blockade, we next evaluated the response of FMOC1 tumors to immune checkpoint inhibitors *in vivo*. FMOC1 cells were inoculated bilaterally into the flanks of FVB/NJ mice by subcutaneous injection and subsequently treated with IgG control, anti-PD-L1 (1.5mg/kg, BioXcell), or anti-CTLA-4 (5mg/kg, BioXcell) via intraperitoneal injection five times over 28 days (n=6-8 tumors per group). FMOC1 tumors exhibited robust sensitivity to both immunotherapies (**Figure 6E and 6G**). At the study endpoint, tumors were harvested and evaluated by IHC for immune cell infiltration. Tumor sections from PD-L1-treated groups showed significantly increased staining of CD8, as well as an elevated CD8/FoxP3 ratio, compared to IgG-treated controls (**Figures 6F**). CTLA-4 tumors exhibited significantly increased staining of CD8 and CD4, as well as an elevated CD8/FoxP3 ratio, compared to IgG-treated controls (**Figures 6H and 6F**). Collectively, these results demonstrated that despite its immunosuppressive TIME, FMOC1 tumors remain responsive to immune checkpoint blockade, accompanied by enhanced T cell infiltration and a shift to a more immunostimulatory landscape.

**Figure 6.**
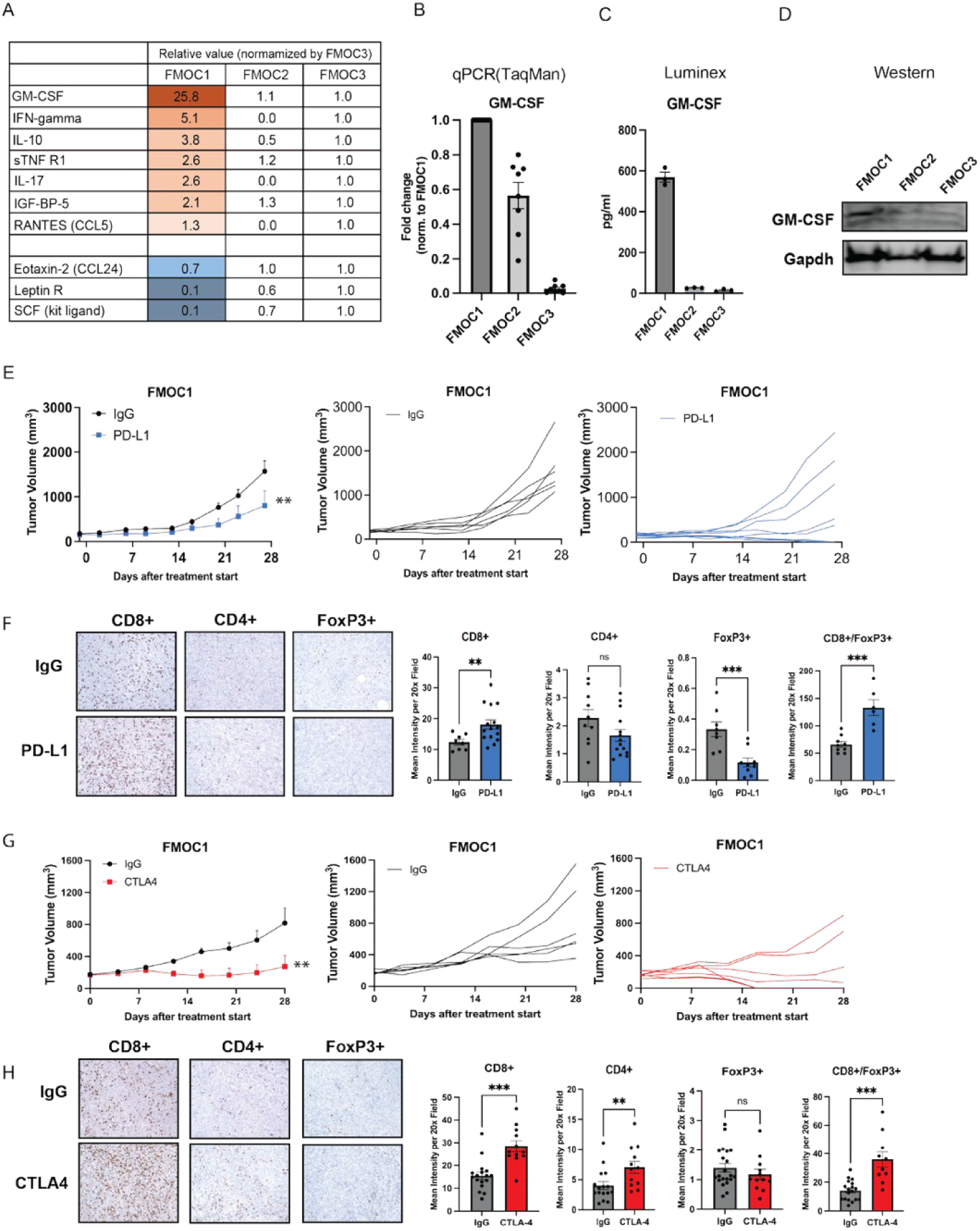
Response of FMOC1 tumors to immune checkpoint inhibitors. (A) The relative value of cytokine expression in FMOC1, FMOC2 and FMOC3 was determined using a cytokine array. Conditioned culture supernatants were collected. The quantitation of protein levels was performed using ImageJ software (Ver. 1.54). (B) RNA was isolated from FMOC cell lines and subjected to qPCR to determine GM-CSF mRNA expression levels in FMOC cell lines. Luminex (C), and immunoblotting (D) were performed to detect GM-CSF protein levels in FMOC cell lines. Gapdh was used as a loading control for immunoblotting. (E-H) FMOC1 tumor are sensitive to immune checkpoint inhibition. (E) FMOC1 tumors were inoculated onto the flanks of FVB/NJ mice. Tumor-bearing mice were treated with either IgG or anti-PD-L1 (1.5mg/kg) therapeutic antibody every 4-5 days for a total of 5 doses. Mean values and SEMs are shown (n=6-8 tumors per group). Spaghetti plots for each tumor presented. (F) Tumor immune infiltrate (CD8, CD4 and FoxP3) was analyzed by IHC and quantified using FIJI version 1.54. Mean values and SEMs are shown (n=8–15 images per group). (G) FMOC1 tumors were inoculated onto the flanks of FVB/NJ mice. Tumor-bearing mice were treated with either IgG, aCTLA-4 (5mg/kg) therapeutic antibody every 4-5 days for a total of 5 doses. Mean values and SEMs are shown (n=6-8 tumors per group). Spaghetti plots for each tumor presented. (H) Tumor immune infiltrate (CD8, CD4 and FoxP3) was analyzed by IHC and quantified using FIJI version 1.54. Mean values and SEMs are shown (n=12–21 images per group). ** P< 0.01; ***P<0.001; ns, not significant.

## Discussion

Despite the clinical success of immune checkpoint inhibitors in some solid tumors, the majority of HNSCC patients fail to achieve durable responses, underscoring the need to better understand mechanisms of immune resistance within the tumor immune microenvironment (TIME). To address this gap, we developed three novel chemically induced syngeneic HNSCC models (FMOC1–3) that exhibit distinct immunogenic and growth characteristics. Preclinical models such as the MOC ^13–16,38^ and 4MOSC ^19^ series, generated through DMBA or 4NQO exposure, respectively, on the C57BL/6 background, have provided important insights into tumor-immune interactions. However, the limited diversity of available models and genetic backgrounds remains a constraint. Here, we introduce the FMOC series, a novel set of DMBA-induced oral cancer cell lines established in the FVB/NJ background. This new platform expands the repository of murine HNSCC models by capturing distinct and functionally relevant tumor-immune phenotypes within a shared genetic background.

The MOC cell line model has been developed and used in many preclinical HNSCC studies^17,18,39,40^. It has been reported that they were generated from different lesions; MOC1 from a mucosal lip lesion, MOC22 from a buccal lesion, and MOC2 from a floor of mouth mass^14^. The FMOC cell line model derived from DMBA-induced tumors developed similar lesions as MOC cell line model (**Figure 1D**). A defining feature of the FMOC series is the marked diversity in mutational profiles across FMOC1, FMOC2, and FMOC3. Despite arising from a common carcinogenic exposure, each line harbors a unique set of oncogenic alterations, including variations in receptor tyrosine kinases (RTKs), PI3K, and TP53 pathway-related gene sets (**Figure 1E and 1F**). This heterogeneity closely mirrors the intertumoral diversity observed in human HNSCC, as well as that reported in the MOC and 4MOSC systems, supporting the biological relevance of these models.

Beyond genetic heterogeneity, the FMOC series exhibits striking differences in tumor-immune interactions. FMOC1 tumors exhibit progressive growth in immunocompetent hosts, enrichment of regulatory T cells (Tregs) and M2 macrophages, and resistance to T cell-mediated cytotoxicity, consistent with an immune-evasive phenotype (**Figure 4**). In contrast, FMOC2 and FMOC3 tumors are characterized by robust infiltration of CD4+ T cells, CD8+ T cells, NK cells and antigen-presenting populations, and undergo spontaneous regression in syngeneic hosts. Functional depletion studies confirm that this tumor control is primarily driven by T cell activity, with cooperative contributions from both CD4+ and CD8+ subsets, particularly in the FMOC3 model (**Figure 5**). These findings position the FMOC series along a spectrum of tumor immune states, with FMOC1 representing an immune-evasive or “cold” tumor, and FMOC2/3 representing immune-inflamed or “hot” tumors that are subject to immune-mediated rejection. While immune heterogeneity has been described in both the MOC and 4MOSC systems, these models were generated under distinct experimental contexts. In contrast, the FMOC series enables direct comparison of immune-evasive and immune-controlled tumor phenotypes derived from a shared carcinogenic exposure, genetic background and experimental pipeline, thereby reducing confounding variables and facilitating the study of tumor-intrinsic mechanisms of immune regulation.

Mechanistically, our data suggests that tumor-intrinsic factors contribute to these divergent immune landscapes. Elevated GM-CSF production by FMOC1 cells, along with enrichment of immunosuppressive myeloid populations, supports a model in which tumor-derived cytokines actively shape a suppressive TIME (**Figure 6**). Although GM-CSF has been shown to have both pro-and anti-tumor effects on the TIME due to its influence on immunosuppressive myeloid populations and dendritic cells, high expression has been associated with poor prognosis in HNSCC^41^. Notably, despite its relatively immunosuppressive baseline state, FMOC1 tumors remain responsive to immune checkpoint blockade. This observation is consistent with prior reports in both MOC and 4MOSC systems demonstrating that even poorly inflamed tumors can still be responsive to immunotherapy.

Importantly, the FMOC models recapitulate key features of HNSCC progression, including orthotopic tumor growth and metastatic potential in both immunocompetent and immunodeficient hosts (**Figure 3**). The differential behavior observed across these settings further highlights the interplay between tumor-intrinsic properties and the TIME in governing tumor progression and metastatic potential. Notably, these properties are most pronounced in FMOC1, consistent with its immune-evasive phenotype and enhanced strong metastatic capacity.

The FMOC models also offer opportunities to interrogate mechanisms of immune priming and memory. Bulk-RNA seq analysis demonstrated increased expression of apoptosis-related pathways in FMOC2 and FMOC3 cells. Consistent with these findings, “educated” CD8^+^ T cells exhibited enhanced cytotoxicity against FMOC2 and FMOC3 cells, suggesting the development of tumor-specific adaptive immunity and highlighting the utility of these models for studying antigen-specific immune responses and immunologic memory in HNSCC (**Figure 1F and 5I**). Because our FMOC cell lines are derived on the FVB/NJ background, they are well suited for use in transgenic mouse systems, including PKCε transgenic models, which have been widely used to study epithelial carcinogenesis^42–46^. As such, FMOC models may facilitate mechanistic studies of skin carcinogenesis and its interaction with the immune microenvironment. One limitation of this model is that, as a DMBA-induced system, the mutational spectrum is dominated by A>T and G>T transversions, which differs from that of HPV-associated human HNSCC^47,48^. Likewise, this mutational spectrum differs from the 4NQO-induced 4MOSC system, which more closely recapitulates tobacco-associated mutagenesis^19^. In addition, tumorigenesis in this model is driven by DMBA, a single chemical carcinogen, whereas human HNSCC develops through many risk factors including complex interaction of tobacco, alcohol, viral infection, and other environmental and genetic factors^49^. Although FMOC2 and FMOC3 regress in immunocompetent mice, this characteristic represents a unique strength of the FMOC model rather than a limitation. Our findings demonstrate that these tumors are controlled through immune-mediated mechanisms, enabling direct comparison between immune-controlled (FMOC2 and FMOC3) and immune-evasive (FMOC1) tumors.

In summary, the FMOC series represents a novel and versatile set of generically related syngeneic HNSCC models that capture key features of tumor heterogeneity and immune diversity. By spanning immune-evasive and immune-controlled states within a unified experimental framework, these models provide a unique platform to investigate the cellular and molecular determinants of tumor-immune interactions and to evaluate therapeutic strategies aimed at overcoming immune resistance in HNSCC.

## Acknowledgments

The authors would like to thank the University of Wisconsin, the Comprehensive Cancer core facilities (Cancer Informatics Shared Resource, Flow Cytometry Laboratory, and the Experimental Animal Pathology Laboratory) supported by P30 CA014520, the Translational Research Initiatives in Pathology (TRIP) laboratory, and the UW Optical Imaging Core. The authors thank the University of Wisconsin-Madison Biotechnology Center Bioinformatics Core Facility (Research Resource Identifier - RRID:SCR_017799) for analysis of WES data.

---

**Supplemental Figure 1.**
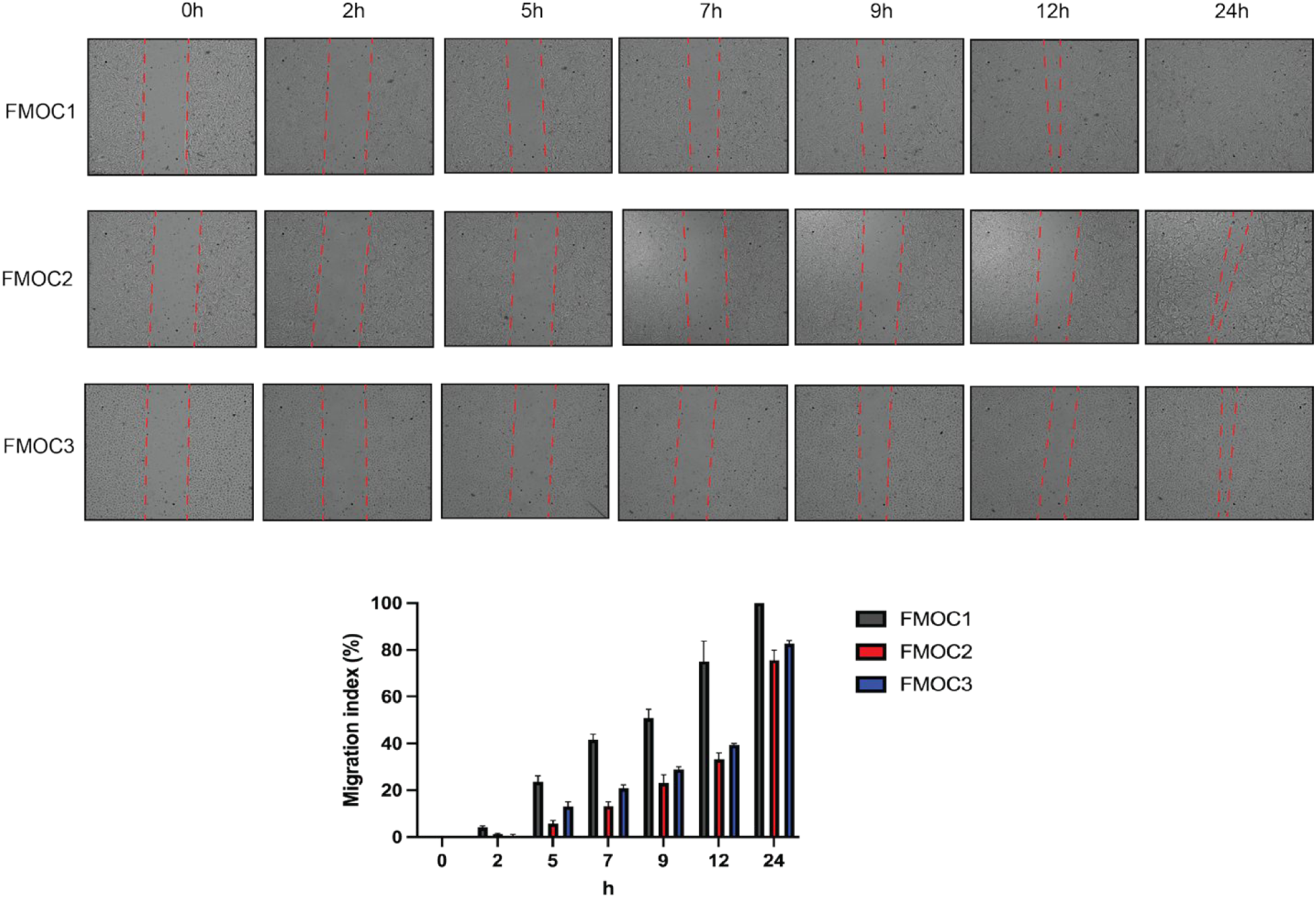
Wound healing assay of FMOC cell lines from 0 to 24 hours incubation following removal of the insert.

**Supplementary Table S1.**

| **Supplementary Table S1. IHC antibodies** |  |  |  |
| --- | --- | --- | --- |
| Antigen | Vendor | Catalog Number | Dilution |
| CD8 | Invitrogen | 14-0808-82 | 1:500 |
| CD4 | Invitrogen | 14-9766-82 | 1:500 |
| NKp46 | Thermo Fisher Scientific, Waltham, MA, USA | PA5-102860 | 1:200 |
| F4/80 | Invitrogen | 14-4801-82 | 1:200 |
| FoxP3 | Invitrogen | 14-5773-826 | 1:200 |
| Ki67 | Cell Signaling Technologies Danvers, MA, USA | 12202 | 1:500 |

**Supplementary Table S2.**

| **Supplementary Table S2. Flow cytometry antibodies** |  |  |  |
| --- | --- | --- | --- |
| Antigen | Fluorescent Marker | Vendor |  |
| CD11b | PE-Vio615 | Miltenyi Biotec | Bergisch Gladbach, Germany |
| CD11c | SuperBright 600 | Thermo Fisher Scientific | Waltham, MA, USA |
| CD170 (Siglec F) | BB515 | BD | Franklin Lakes, NJ, USA |
| CD25 | BrilliantViolet 510 | Biolegend | San Diego, CA, USA |
| CD3 | AlexaFluor 488 | Biolegend | San Diego, CA, USA |
| CD4 | PerCP-eFluor 710 | Thermo Fisher Scientific | Waltham, MA, USA |
| CD19 | SB 702 | Thermo Fisher Scientific | Waltham, MA, USA |
| CD27 | PE-Vio615 | Miltenyi Biotec | Bergisch Gladbach, Germany |
| CD45 | PE-Cy7 | Biolegend | San Diego, CA, USA |
| CD69 | PE-Cy5 | Biolegend | San Diego, CA, USA |
| CD80 | SuperBright 436 | Thermo Fisher Scientific | Waltham, MA, USA |
| CD8a | AlexaFluor 700 | Biolegend | San Diego, CA, USA |
| CD206 | APC | Biolegend | San Diego, CA, USA |
| F4/80 | SuperBright 702 | Thermo Fisher Scientific | Waltham, MA, USA |
| Live/Dead | GhostRed 780 | Tonbo Biosciences | San Diego, CA, USA |
| Ly6G | PE | Tonbo Biosciences | San Diego, CA, USA |
| MHC-II (I-A/I-E) | BrilliantViolet 510 | BD | Franklin Lakes, NJ, USA |
| NKp46 | SuperBright 436 | Thermo Fisher Scientific | Waltham, MA, USA |
| FoxP3 | PE | Tonbo Biosciences | San Diego, CA, USA |
| PD1 | SB 600 | Thermo Fisher Scientific | Waltham, MA, USA |
| Perforin | APC | Biolegend | San Diego, CA, USA |
| FcεR1a | PerCP-e710 | Thermo Fisher Scientific | Waltham, MA, USA |

**Supplementary Table S3.**
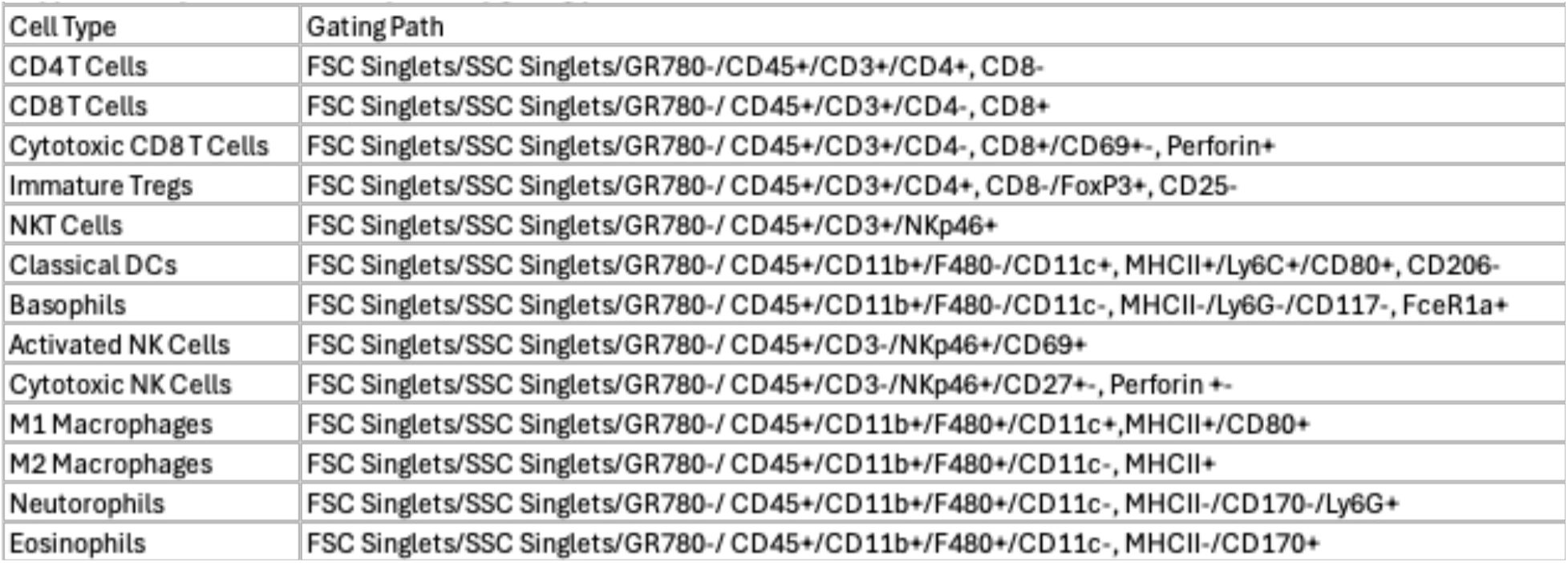

**Supplementary Table S4.**
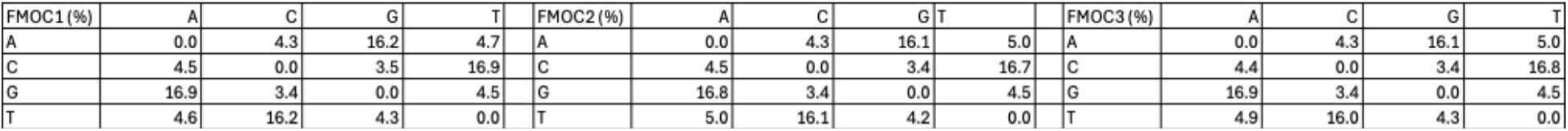

**Supplementary Table S5.** Bulk RNA-Seq data

## Notes

### Competing Interest Statement

The authors have declared no competing interest.

## References

1. Siegel, R.L., Kratzer, T.B., Giaquinto, A.N., Sung, H., and Jemal, A. (2025). Cancer statistics, 2025. CA: a cancer J. Clin. 75, 10–45. 10.3322/caac.21871.

2. Zhou, G., Liu, Z., and Myers, J.N. (2016). TP53 Mutations in Head and Neck Squamous Cell Carcinoma and Their Impact on Disease Progression and Treatment Response. J. Cell. Biochem. 117, 2682–2692. 10.1002/jcb.25592.

3. Stransky, N., Egloff, A.M., Tward, A.D., Kostic, A.D., Cibulskis, K., Sivachenko, A., Kryukov, G.V., Lawrence, M., Sougnez, C., McKenna, A., et al. (2011). The Mutational Landscape of Head and Neck Squamous Cell Carcinoma. Science (New York, NY). 10.1126/science.1208130.

4. Pickering, C.R., Zhang, J., Yoo, S.Y., Bengtsson, L., Moorthy, S., Neskey, D.M., Zhao, M., Alves, M.V.O., Chang, K., Drummond, J., et al. (2013). Integrative Genomic Characterization of Oral Squamous Cell Carcinoma Identifies Frequent Somatic Drivers. Cancer Discov. 3, 770–781. 10.1158/2159-8290.cd-12-0537.

5. Heinolainen, A., Nguyen, B., Silén, S., Renkonen, R., and Koskinen, M. (2025). Survival and data-driven phenotypes in head and neck cancer. Sci. Rep. 15, 5985. 10.1038/s41598-025-89053-6.

6. L., F.R., Jr., B.G., Jerome, F., Joel, G., Dimitrios, C.A., Lisa, L., Kevin, H., Stefan, K., E., V.E., Caroline, E., et al. (2016). Nivolumab for Recurrent Squamous-Cell Carcinoma of the Head and Neck. N. Engl. J. Med. 375, 1856–1867. 10.1056/nejmoa1602252.

7. Uppaluri, R., Haddad, R.I., Tao, Y., Tourneau, C.L., Lee, N.Y., Westra, W., Chernock, R., Tahara, M., Harrington, K.J., Klochikhin, A.L., et al. (2025). Neoadjuvant and Adjuvant Pembrolizumab in Locally Advanced Head and Neck Cancer. N. Engl. J. Med. 393, 37–50. 10.1056/nejmoa2415434.

8. Seiwert, T.Y., Burtness, B., Mehra, R., Weiss, J., Berger, R., Eder, J.P., Heath, K., McClanahan, T., Lunceford, J., Gause, C., et al. (2016). Safety and clinical activity of pembrolizumab for treatment of recurrent or metastatic squamous cell carcinoma of the head and neck (KEYNOTE-012): an open-label, multicentre, phase 1b trial. Lancet Oncol. 17, 956–965. 10.1016/s1470-2045(16)30066-3.

9. Burtness, B., Harrington, K.J., Greil, R., Soulières, D., Tahara, M., Castro, G. de, Psyrri, A., Basté, N., Neupane, P., Bratland, Å., et al. (2019). Pembrolizumab alone or with chemotherapy versus cetuximab with chemotherapy for recurrent or metastatic squamous cell carcinoma of the head and neck (KEYNOTE-048): a randomised, open-label, phase 3 study. Lancet 394, 1915–1928. 10.1016/s0140-6736(19)32591-7.

10. Larkins, E., Blumenthal, G.M., Yuan, W., He, K., Sridhara, R., Subramaniam, S., Zhao, H., Liu, C., Yu, J., Goldberg, K.B., et al. (2017). FDA Approval Summary: Pembrolizumab for the Treatment of Recurrent or Metastatic Head and Neck Squamous Cell Carcinoma with Disease Progression on or After Platinum-Containing Chemotherapy. Oncol. 22, 873–878. 10.1634/theoncologist.2016-0496.

11. Elmusrati, A., Wang, J., and Wang, C.-Y. (2021). Tumor microenvironment and immune evasion in head and neck squamous cell carcinoma. Int. J. Oral Sci. 13, 24. 10.1038/s41368-021-00131-7.

12. Tsai, C.-C., Hsu, Y.-C., Chu, T.-Y., Hsu, P.-C., and Kuo, C.-Y. (2025). Immune Evasion in Head and Neck Squamous Cell Carcinoma: Roles of Cancer-Associated Fibroblasts, Immune Checkpoints, and TP53 Mutations in the Tumor Microenvironment. Cancers 17, 2590. 10.3390/cancers17152590.

13. Ku, T.K.S., Nguyen, D.C., Karaman, M., Gill, P., Hacia, J.G., and Crowe, D.L. (2007). Loss of p53 Expression Correlates with Metastatic Phenotype and Transcriptional Profile in a New Mouse Model of Head and Neck Cancer. Molecular Cancer Research 5, 351–362. 10.1158/1541-7786.mcr-06-0238.

14. Judd, N.P., Winkler, A.E., Murillo-Sauca, O., Brotman, J.J., Law, J.H., Lewis, J.S., Dunn, G.P., Bui, J.D., Sunwoo, J.B., and Uppaluri, R. (2012). ERK1/2 regulation of CD44 modulates oral cancer aggressiveness. Cancer Research 72, 365–374. 10.1158/0008-5472.can-11-1831.

15. Onken, M.D., Winkler, A.E., Kanchi, K.-L., Chalivendra, V., Law, J.H., Rickert, C.G., Kallogjeri, D., Judd, N.P., Dunn, G.P., Piccirillo, J.F., et al. (2014). A surprising cross-species conservation in the genomic landscape of mouse and human oral cancer identifies a transcriptional signature predicting metastatic disease. Clin Cancer Res Official J Am Assoc Cancer Res 20, 2873–2884. 10.1158/1078-0432.ccr-14-0205.

16. Kono, M., Saito, S., Egloff, A.M., Allen, C.T., and Uppaluri, R. (2022). The mouse oral carcinoma (MOC) model: A 10-year retrospective on model development and head and neck cancer investigations. Oral Oncol. 132, 106012. 10.1016/j.oraloncology.2022.106012.

17. Kostecki, K.L., Harmon, R.L., Iida, M., Harris, M.A., Crossman, B.E., Bruce, J.Y., Salgia, R., and Wheeler, D.L. (2025). Axl Regulation of NK Cell Activity Creates an Immunosuppressive Tumor Immune Microenvironment in Head and Neck Cancer. Cancers 17, 994. 10.3390/cancers17060994.

18. Kostecki, K.L., Iida, M., Wiley, A.L., Kimani, S., Mehall, B., Tetreault, K., Alexandridis, R., Yu, M., Hong, S., Salgia, R., et al. (2022). Dual Axl/MerTK inhibitor INCB081776 creates a proinflammatory tumor immune microenvironment and enhances anti-PDL1 efficacy in head and neck cancer. Head neck 45, 1255–1271. 10.1002/hed.27340.

19. Wang, Z., Wu, V.H., Allevato, M.M., Gilardi, M., He, Y., Callejas-Valera, J.L., Vitale-Cross, L., Martin, D., Amornphimoltham, P., Mcdermott, J., et al. (2019). Syngeneic animal models of tobacco-associated oral cancer reveal the activity of in situ anti-CTLA-4. Nat Commun 10, 5546. 10.1038/s41467-019-13471-0.

20. Mao, L., Zhou, J.-J., Xiao, Y., Yang, Q.-C., Yang, S.-C., Wang, S., Wu, Z.-Z., Xiong, H.-G., Yu, H.-J., and Sun, Z.-J. (2023). Immunogenic hypofractionated radiotherapy sensitising head and neck squamous cell carcinoma to anti-PD-L1 therapy in MDSC-dependent manner. Br. J. Cancer 128, 2126–2139. 10.1038/s41416-023-02230-0.

21. Suit, H.D., Sedlacek, R.S., and Zietman, A. (1988). Quantitative transplantation assays of spontaneous tumors of the C3H mouse as allografts in athymic NCr/Sed-nu/nu nude mice and isografts in C3Hf/Sed mice. Cancer Res 48, 4525–4528.

22. Iida, M., Crossman, B.E., Kostecki, K.L., Glitchev, C.E., Kranjac, C.A., Crow, M.T., Adams, J.M., Liu, P., Ong, I., Yang, D.T., et al. (2024). MerTK Drives Proliferation and Metastatic Potential in Triple-Negative Breast Cancer. Int. J. Mol. Sci. 25, 5109. 10.3390/ijms25105109.

23. McDaniel, N.K., Iida, M., Nickel, K.P., Longhurst, C.A., Fischbach, S.R., Rodems, T.S., Kranjac, C.A., Bo, A.Y., Luo, Q., Gallagher, M.M., et al. (2019). AXL Mediates Cetuximab and Radiation Resistance Through Tyrosine 821 and the c-ABL Kinase Pathway in Head and Neck Cancer. Clin. cancer Res. : Off. J. Am. Assoc. Cancer Res. 26, 4349–4359. 10.1158/1078-0432.ccr-19-3142.

24. Iida, M., McDaniel, N.K., Kostecki, K.L., Welke, N.B., Kranjac, C.A., Liu, P., Longhurst, C., Bruce, J.Y., Hong, S., Salgia, R., et al. (2022). AXL regulates neuregulin1 expression leading to cetuximab resistance in head and neck cancer. Bmc Cancer 22, 447. 10.1186/s12885-022-09511-6.

25. McDaniel, N.K., Cummings, C.T., Iida, M., Hülse, J., Pearson, H.E., Vasileiadi, E., Parker, R.E., Orbuch, R.A., Ondracek, O.J., Welke, N.B., et al. (2018). MERTK Mediates Intrinsic and Adaptive Resistance to AXL-targeting Agents. Mol. cancer Ther. 17, 2297–2308. 10.1158/1535-7163.mct-17-1239.

26. Pearson, H.E., Iida, M., Orbuch, R.A., McDaniel, N.K., Nickel, K.P., Kimple, R.J., Arbiser, J., and Wheeler, D.L. (2017). Overcoming resistance to cetuximab with honokiol, a small-molecule polyphenol. Mol Cancer Ther 17, molcanther.0384.2017. 10.1158/1535-7163.mct-17-0384.

27. Crossman, B.E., Harmon, R.L., Iida, M., Adams, J.M., Lin, C.Y., Glitchev, C.E., Juang, T.D., Kerr, S.C., Alexandridis, R.A., Hyun, M., et al. (2025). Tumor-associated MerTK promotes a pro-inflammatory microenvironment and enhances immune checkpoint inhibitor response in triple-negative breast cancer. Front. Oncol. 15, 1579214. 10.3389/fonc.2025.1579214.

28. Chen, S., Zhou, Y., Chen, Y., and Gu, J. (2018). fastp: an ultra-fast all-in-one FASTQ preprocessor. Bioinformatics 34, i884–i890. 10.1093/bioinformatics/bty560.

29. Dobin, A., Davis, C.A., Schlesinger, F., Drenkow, J., Zaleski, C., Jha, S., Batut, P., Chaisson, M., and Gingeras, T.R. (2012). STAR: ultrafast universal RNA-seq aligner. Bioinformatics 29, 15–21. 10.1093/bioinformatics/bts635.

30. Frankish, A., Diekhans, M., Ferreira, A.-M., Johnson, R., Jungreis, I., Loveland, J., Mudge, J.M., Sisu, C., Wright, J., Armstrong, J., et al. (2019). GENCODE reference annotation for the human and mouse genomes. Nucleic Acids Res. 47, D766–D773. 10.1093/nar/gky955.

31. Liao, Y., Smyth, G.K., and Shi, W. (2014). featureCounts: an efficient general purpose program for assigning sequence reads to genomic features. Bioinformatics 30, 923–930. 10.1093/bioinformatics/btt656.

32. Love, M.I., Huber, W., and Anders, S. (2014). Moderated estimation of fold change and dispersion for RNA-seq data with DESeq2. Genome Biol. 15, 550. 10.1186/s13059-014-0550-8.

33. Kolde, R. (2010). pheatmap: Pretty Heatmaps. CRAN: Contrib. Packag. 10.32614/cran.package.pheatmap.

34. Wickham, H. (2016). ggplot2, Elegant Graphics for Data Analysis. Use R! 10.1007/978-3-319-24277-4.

35. Yan, L. (2021). ggvenn: Draw Venn Diagram by “ggplot2.” CRAN: Contrib. Packag. 10.32614/cran.package.ggvenn.

36. Gu, Z., Gu, L., Eils, R., Schlesner, M., and Brors, B. (2014). circlize Implements and enhances circular visualization in R. Bioinform. (Oxf., Engl.) 30, 2811–2812. 10.1093/bioinformatics/btu393.

37. Gu, Z., Eils, R., and Schlesner, M. (2016). Complex heatmaps reveal patterns and correlations in multidimensional genomic data. Bioinformatics 32, 2847–2849. 10.1093/bioinformatics/btw313.

38. Zolkind, P., and Uppaluri, R. (2017). Checkpoint immunotherapy in head and neck cancers. 1–15. 10.1007/s10555-017-9694-9.

39. Jin, W.J., Jagodinsky, J.C., Vera, J.M., Clark, P.A., Zuleger, C.L., Erbe, A.K., Ong, I.M., Le, T., Tetreault, K., Berg, T., et al. (2023). NK cells propagate T cell immunity following in situ tumor vaccination. Cell Rep. 42, 113556. 10.1016/j.celrep.2023.113556.

40. Jin, W.J., Erbe, A.K., Schwarz, C.N., Jaquish, A.A., Anderson, B.R., Sriramaneni, R.N., Jagodinsky, J.C., Bates, A.M., Clark, P.A., Le, T., et al. (2020). Tumor-Specific Antibody, Cetuximab, Enhances the In Situ Vaccine Effect of Radiation in Immunologically Cold Head and Neck Squamous Cell Carcinoma. Front. Immunol. 11, 591139. 10.3389/fimmu.2020.591139.

41. Lopes-Santos, G., Tjioe, K.C., Magalhaes, M.A. de O., and Oliveira, D.T. (2023). The role of granulocyte-macrophage colony-stimulating factor in head and neck cancer. Arch. Oral Biol. 147, 105641. 10.1016/j.archoralbio.2023.105641.

42. Wheeler, D.L., Reddig, P.J., Ness, K.J., Leith, C.P., Oberley, T.D., and Verma, A.K. (2005). Overexpression of Protein Kinase C-ε in the Mouse Epidermis Leads to a Spontaneous Myeloproliferative-Like Disease. Am. J. Pathol. 166, 117–126. 10.1016/s0002-9440(10)62237-7.

43. Wheeler, D.L., Martin, K.E., Ness, K.J., Li, Y., Dreckschmidt, N.E., Wartman, M., Ananthaswamy, H.N., Mitchell, D.L., and Verma, A.K. (2004). Protein kinase C epsilon is an endogenous photosensitizer that enhances ultraviolet radiation-induced cutaneous damage and development of squamous cell carcinomas. Cancer Research 64, 7756–7765. 10.1158/0008-5472.can-04-1881.

44. Singh, A., Singh, A., Bauer, S.J., Wheeler, D.L., Havighurst, T.C., Kim, K., and Verma, A.K. (2015). Genetic deletion of TNFα inhibits ultraviolet radiation-induced development of cutaneous squamous cell carcinomas in PKCε transgenic mice via inhibition of cell survival signals. Carcinogenesis. 10.1093/carcin/bgv162.

45. Li, Y., Wheeler, D., Alters, W., Chaiswing, L., Verma, A., and Oberley, T. (2005). Early Epidermal Destruction with Subsequent Epidermal Hyperplasia Is a Unique Feature of the Papilloma-Independent Squamous Cell Carcinoma Phenotype in PKCε Overexpressing Transgenic Mice. Toxicologic Pathology 33, 684–694. 10.1080/01926230500323441.

46. Taketo, M., Schroeder, A.C., Mobraaten, L.E., Gunning, K.B., Hanten, G., Fox, R.R., Roderick, T.H., Stewart, C.L., Lilly, F., and Hansen, C.T. (1991). FVB/N: an inbred mouse strain preferable for transgenic analyses. Proc. Natl. Acad. Sci. 88, 2065–2069. 10.1073/pnas.88.6.2065.

47. Sabatini, M.E., and Chiocca, S. (2020). Human papillomavirus as a driver of head and neck cancers. Br. J. Cancer 122, 306–314. 10.1038/s41416-019-0602-7.

48. Gendreizig, S., Martínez-Ruiz, L., López-Rodríguez, A., Pabla, H., Hose, L., Brasch, F., Busche, T., Escames, G., Sudhoff, H., Scholtz, L.U., et al. (2024). Human papillomavirus-associated head and neck squamous cell carcinoma cells lose viability during triggered myocyte lineage differentiation. Cell Death Dis. 15, 517. 10.1038/s41419-024-06867-4.

49. Johnson, D.E., Burtness, B., Leemans, C.R., Lui, V.W.Y., Bauman, J.E., and Grandis, J.R. (2020). Head and neck squamous cell carcinoma. Nat. Rev. Dis. Prim. 6, 92. 10.1038/s41572-020-00224-3.

